# Coupled intracellular redox and extracellular respiration sensing for quantitative oxidative stress profiling

**DOI:** 10.64898/2026.05.30.727573

**Authors:** Shirin Parvin, Deon Ploessl, Zheyuan Zhang, Zengyi Shao, Meng Lu

**Affiliations:** Department of Electrical and Computer Engineering, Iowa State University, Ames, Iowa 50011, USA; Department of Chemical and Biological Engineering, Iowa State University, Ames, Iowa 50011, USA; Department of Mechanical Engineering, Iowa State University, Ames, Iowa 50011, USA

**Keywords:** Reactive oxygen species, genetically encoded sensor, dissolved oxygen sensor, biomanufacturing

## Abstract

Quantitative assessment of cellular oxidative stress requires simultaneous measurement of intracellular redox state and extracellular respiratory activity, yet integrated sensing approaches remain limited. Here, we present a dual-fluorescent sensing platform combining a genetically encoded redox biosensor (roGFP2□Tsa2ΔC_R_) with an optical oxygen sensor embedded in microwell plates for parallel, non-invasive quantification of intracellular reactive oxygen species (ROS) and oxygen consumption rates (OCR) in industrial yeast systems. The roGFP2-based sensor was stably expressed in *Saccharomyces cerevisiae* (*S. cerevisiae*) and *Yarrowia lipolytica* (*Y. lipolytica*), enabling dynamic monitoring of oxidative stress at population, single-cell, and subcellular levels, while oxygen-sensitive films provided real-time respiration measurements. Using this platform, we identified distinct redox–respiration phenotypes between the two yeasts. Crabtree-positive *S. cerevisiae* exhibited low OCR and mitochondrial ROS during glucose cultivation, whereas growth on glycerol increased OCR and mitochondrial ROS by ~2.5-fold and 12%, respectively. In contrast, the obligate respiratory yeast *Y. lipolytica* displayed 3-fold higher OCR and 16% lower mitochondrial ROS than respiring *S. cerevisiae*, indicating differences in respiratory oxidative burden. Antimycin A treatment reduced OCR by 60% in respiring *S. cerevisiae* while increasing mitochondrial ROS by 35%, whereas *Y. lipolytica* showed greater resistance to respiratory and oxidative perturbations. By integrating intracellular redox sensing with extracellular oxygen measurements, this platform enables quantitative coupling of redox state and respiration in living cells. The approach provides a scalable framework for evaluating cellular fitness, stress tolerance, and metabolic state in biomanufacturing and synthetic biology.

## 1. Introduction

Microbial cell factories, including conventional and non-conventional hosts, are increasingly used in biomanufacturing because of their metabolic flexibility and engineering potential. In these systems, cell fitness, including stress response, metabolic activity, and redox balance, is critical to production efficiency (Wang et al., 2024). Stressors, such as oxidative stress, nutrient limitation, and byproduct accumulation can disrupt cellular processes, causing reduced growth, metabolic imbalance, and genetic instability (Sinharoy et al., 2021; Chevallier et al., 2020). Understanding the relationship between oxidative stress, antioxidant capacity, and metabolism is therefore important for improving process robustness and productivity (Phaneuf et al., 2024; Liu et al., 2023). Reactive oxygen species (ROS), primarily generated during mitochondrial oxidative phosphorylation, play important roles in signaling and metabolism but can damage proteins, lipids, and DNA when accumulated excessively (Mittler; 2017; Zorov et al., 2014). During respiration, electrons flow through the mitochondrial electron transport chain (ETC) to drive ATP synthesis, while electron leakage from complexes I and III generates superoxide and downstream ROS species, as illustrated in **Fig. 1(a)**.

**Fig. 1.**
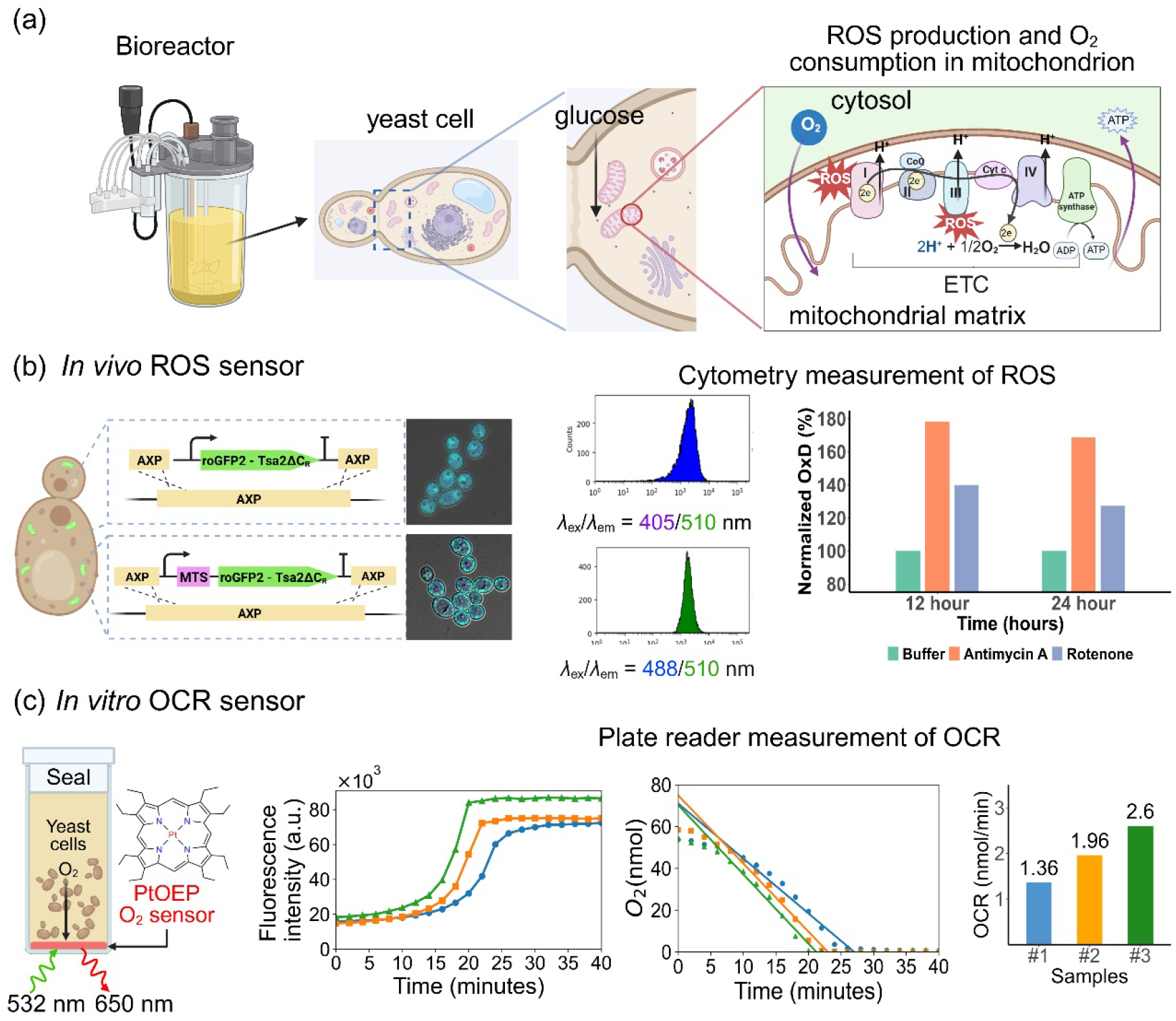
Mitochondrial electron transport chain linking ROS production and oxygen respiration. (a) Schematic of mitochondrial respiration and ROS generation. Electrons flow through the electron transport chain to reduce oxygen to water and generate ATP, while electron leakage generates ROS, which contributes to oxidative stress. (b-c) An integrated fluorescent sensing platform combining a (b) genetically encoded in vivo ROS sensor enables spatially resolved intracellular ROS monitoring under different chemical perturbations, and a (c) micro-well embedded optical oxygen sensor for extracellular respiration measurement.

Current ROS assays commonly rely on fluorescent dyes and often suffer from limited specificity or dose sensitivity (Eruslanov and Kusmartsev, 2010). In contrast, genetically encoded roGFP-based sensors are ratiometric and enable real-time quantification of intracellular oxidative stress at both population and single-cell levels without disrupting normal cellular function (Kauffman et al., 2016; Kuznesov et al., 2011; Wojtala et al., 2014; Morgan et al., 2011; Meyer and Dick, 2010). For respiration measurements, Clark-type electrodes remain a standard method (Li and Graham, 2012; Clark et al., 1958), while oxygen-sensitive fluorescent dyes provide a high-throughput alternative for monitoring oxygen uptake (Wolfbeis, 2015; Melnikov et al., 2022). Combined measurement of ROS and respiration enables direct correlation of oxidative stress with mitochondrial activity, offering a more comprehensive assessment of cellular fitness and redox homeostasis during metabolic transitions.

Here, we present an integrated sensing platform combining a genetically encoded redox biosensor (roGFP2□Tsa2ΔC_R_) for intracellular ROS measurement with an extracellular optical oxygen sensor for respiration monitoring, as shown in **Fig. 1(b)**. Using *Saccharomyces cerevisiae* (*S. cerevisiae*) and *Yarrowia lipolytica* (*Y. lipolytica*), we quantified dynamic changes in intracellular redox state and oxygen consumption during batch cultivation. To the best of our knowledge, this study provides the first characterization of roGFP2 performance in *Y. lipolytica*. The integrated measurements revealed dose-dependent OCR suppression and compartment-specific ROS accumulation under both chemical and metabolic perturbations. Compared with glucose-grown cells, glycerol-grown *S. cerevisiae* exhibited greater respiratory inhibition and but lower oxidative stress, whereas *Y. lipolytica* maintained more stable respiration and redox balance. Together, the combined OCR and roGFP measurements establish a quantitative link between ETC activity, metabolic state, and intracellular redox dynamics, demonstrating the utility of dual-sensing platforms for evaluating cellular fitness in microbial biomanufacturing.

## 2. Materials and Methods

### 2.1 Materials and reagents

Laboratory yeast strains *S. cerevisiae* BY4741 (ATCC 201388) and *Y. lipolytica* po1f (ATCC MYA-2613) were obtained from ATCC. PCR amplification and DNA assembly were performed using Q5® High-Fidelity Master Mix and the NEBuilder® HiFi DNA Assembly Kit (New England Biolabs), respectively. *E. coli* NEB 10-beta competent cells were used for plasmid transformation, and plasmids were purified using the QIAprep Spin Miniprep Kit (Qiagen). Restriction digests were carried out using FastDigest™ enzymes (Thermo Fisher Scientific). For dissolved oxygen sensor fabrication, PDMS (Sylgard® 184), platinum (II) octaethylporphyrin (PtOEP), ethanol, and dimethyl sulfoxide (DMSO) were purchased from Sigma-Aldrich. Antimycin A (AA), rotenone, furfural, glucose, acetic acid, dithiothreitol (DTT), diamide, sorbitol, NaCl, Tris-HCl, hydrogen peroxide, LB medium, yeast extract, peptone, glycerol, and Phosphate Buffered Saline (PBS) were obtained from Sigma-Aldrich or Fisher Scientific.

### 2.2 Cloning of roGFP2-Tsa2ΔC_R_ sensor in yeast strains

The H□O□ sensor plasmid p415 TEF Su9-roGFP2□Tsa2ΔC_R_ was a gift from Tobias Dick (Addgene #83240) (Morgan et al., 2016). The roGFP2-Tsa2ΔC_R_ coding sequence and mitochondrial targeting sequence (MTS) were codon optimized for expression in *Y. lipolytica* and synthesized by GenScript (Piscataway, NJ). Promoters, terminators, ORFs, and homologous arms were amplified by PCR and assembled into genomic integration cassettes containing the roGFP2□Tsa2ΔC_R_ sensor and Ura3 selection marker using HiFi assembly. The constructs were integrated into the *S. cerevisiae* IDH1 locus and *Y. lipolytica* AXP locus following the LINEAR integration method (Ploessl et al., 2022). Transformants were selected on SC-Ura agar plates at 30 °C and verified by genomic PCR. Strains containing roGFP2-Tsa2ΔC_R_ with or without the MTS are referred to as mitochondrial and cytosolic roGFP2-Tsa2ΔC_R_ strains, respectively.

### 2.3. Cell cultures and sampling conditions

Mitochondrial and cytosolic roGFP2-Tsa2ΔC_R_ strains of *S. cerevisiae* and *Y. lipolytica* were cultured in 50 mL YPD or YPG medium in 250 mL shaking flasks at 30 °C and 250 rpm. YPD contained 2% glucose, while YPG contained 2% ethanol and 1% glycerol, supplemented with 10 g/L yeast extract and 20 g/L peptone. Cultures were inoculated to an initial OD_600_ of 0.1, and growth was monitored using a UV-Vis spectrophotometer (Cary 60, Agilent). Cells were sampled during cultivation for OCR and ROS measurements, washed twice with isosmotic buffer (0.1 M sorbitol, 0.1 M NaCl, 0.1 M Tris-HCl, pH 7.4) and resuspended to OD□□□ = 0.2.

### 2.4. Measurement of roGFP2-Tsa2ΔC_R_ sensor

Flow cytometry was used to evaluate cellular heterogeneity of roGFP2-Tsa2ΔC_R_ sensor strains under oxidative stress. Cultured cells were washed twice with isosmotic buffer, resuspended to OD□□□ = 0.2, and incubated with chemical agents in 96-well plates at 30 °C for 30 min at 250 rpm. After treatment, 100 μL of each sample was diluted with 200 μL PBS and analyzed using a BD FACSDiscover A8 flow cytometer with 405 nm and 488 nm excitation and 515 nm emission detection. Subcellular localization and redox responses were visualized using a Leica Stellaris 5 confocal microscope with a 63× oil immersion objective. Fluorescence images were acquired using 405 nm and 488 nm excitation with emission collected at 515 nm. Cells were immobilized on #1.0 glass coverslips using agarose pads, and multiple fields of view were collected for each sample. Images were processed using ImageJ software.

### 2.5. Calculation of cell oxidative stress

Raw fluorescence data were processed to obtain roGFP2 emission intensities at 515 nm under 405 nm and 488 nm excitation, denoted as *I*_405_ and *I*_488_, respectively. The fluorescence ratio R = *I*_405_/*I*_488_ is commonly used to quantify roGFP redox state because it minimizes variability arising from probe expression levels (Morgan et al., 2011; Radzinski et al., 2018; Chandrasekharan et al., 2019; Ugalde et al., 2022). However, this ratio remains sensitive to instrumental settings and measurement conditions. Therefore, the degree of oxidation (OxD), representing the fraction of oxidized roGFP molecules, was used for quantitative comparison across measurement modalities. OxD was calculated using the following equation:

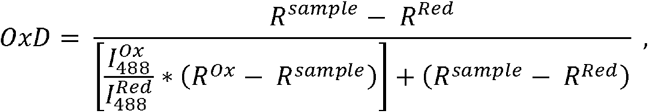

where 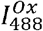 and 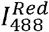 represent the fluorescence intensities at 488 nm under fully oxidized (diamide-treated) and fully reduced (DTT-treated) conditions, respectively (Morgan et al., 2016; Calabrese et al., 2019; Pastor-Flores et al., 2017). OxD values for individual cells were calculated using the median fluorescence intensities of DTT- and diamide-treated populations as OxD references of 0 and 1, respectively. Statistical metrics, including medians, interquartile ranges, means, standard deviations, and correlation coefficients, were computed using custom Python scripts.

### 2.6. Preparation of microwell plate for dissolved oxygen sensing

To assess cellular respiration, oxygen-sensitive microwell plates were fabricated by coating each well with a PtOEP-doped PDMS film, as previously described (Penso et al., 2021). PtOEP was dissolved in uncured PDMS at 0.1% w/w and degassed before deposition. Microwell plates were oxygen plasma-treated (100 W, 1 min) to improve adhesion, followed by dispensing 20 μL of the PtOEP-PDMS mixture into each well and centrifugation at 500 rpm for 30 s to form a uniform film. The plates were cured at 60 °C for 2 h, rinsed with 70% ethanol, air dried, and vacuum-sealed until use.

### 2.7. Measurement of cell respiration rate

PtOEP fluorescence intensity is inversely proportional to dissolved oxygen concentration. Fluorescence measurements were performed using a microplate reader (Synergy H1, Agilent) with excitation at 532 nm and emission at 650 nm ± 20 nm. For respiration assays, cells were harvested, washed, and resuspended in fresh medium to OD□□□ = 0.2. After adding 200 μL of cell suspension and chemical reagents, microwells were sealed with soft paraffin to minimize gas exchange and monitored at 30 °C for 60 min with 2 min intervals. Raw fluorescence intensities were converted to dissolved oxygen concentrations using the Stern-Volmer equation:

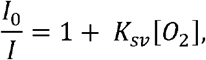

where *I*_0_ is the fluorescence intensity in fully deoxygenated media, *I* is the intensity at oxygen concentration [O_2_], and *K*_sv_ is the Stern-Volmer constant (Boaz and Rollefson, 1950; Keizer, 1983; Gehlen, 2020). The Stern-Volmer equation relates fluorescence quenching to oxygen levels through inversion of the normalized intensity signal, which is then used as a measure of mitochondrial function (Will et al., 2006).To account for well-to-well variability, each microwell was calibrated using oxygen-saturated media (~0.26 mM dissolved oxygen) and fully deoxygenated media. Deoxygenated media contained 80 mM sodium sulfite with cobalt chloride catalyst to scavenge oxygen (Ghaly and Kok, 1988; Miron, 1979; Snavely, 1971). OCR was determined by linear regression over a dynamic window, with the slope corresponding to the oxygen consumption rate. Because *Y. lipolytica* exhibits dimorphic growth unlike *S. cerevisiae*, OCR values were normalized to OD□□□ for cross-species comparison.

## 3. Results and discussions

### 3.1. Characterization of intracellular ROS levels using roGFP

The yeast strains expressing cytosolic and mitochondrial roGFP2-Tsa2ΔC_R_ were characterized for fluorescence response and subcellular localization. **Fig. 2(b)** shows the AlphaFold3-predicted structure of roGFP2-Tsa2ΔC_R_, in which the peroxiredoxin Tsa2ΔC_R_ is fused to roGFP2 through a GS linker. Tsa2ΔC_R_ functions as a hydrogen peroxide-sensitive relay that modulates the redox state of roGFP2, resulting in fluorescence changes under oxidized and reduced conditions. Fluorescence characterization was performed using a microplate reader (Synergy H1, Agilent). Cells were pelleted in 96-well plates, and emission at 515 nm was recorded while scanning excitation wavelengths from 375-488 nm. As shown in **Fig. 2(c)**, roGFP2 exhibited two excitation peaks at 405 nm and 488 nm, corresponding to oxidized and reduced states, respectively. Oxidation increases the 405 nm excitation peak through disulfide bond formation, whereas reduction enhances the 488 nm peak. Therefore, the fluorescence ratio (*I*_405_/*I*_488_) provides a quantitative measure of intracellular redox state.

**Figure 2.**
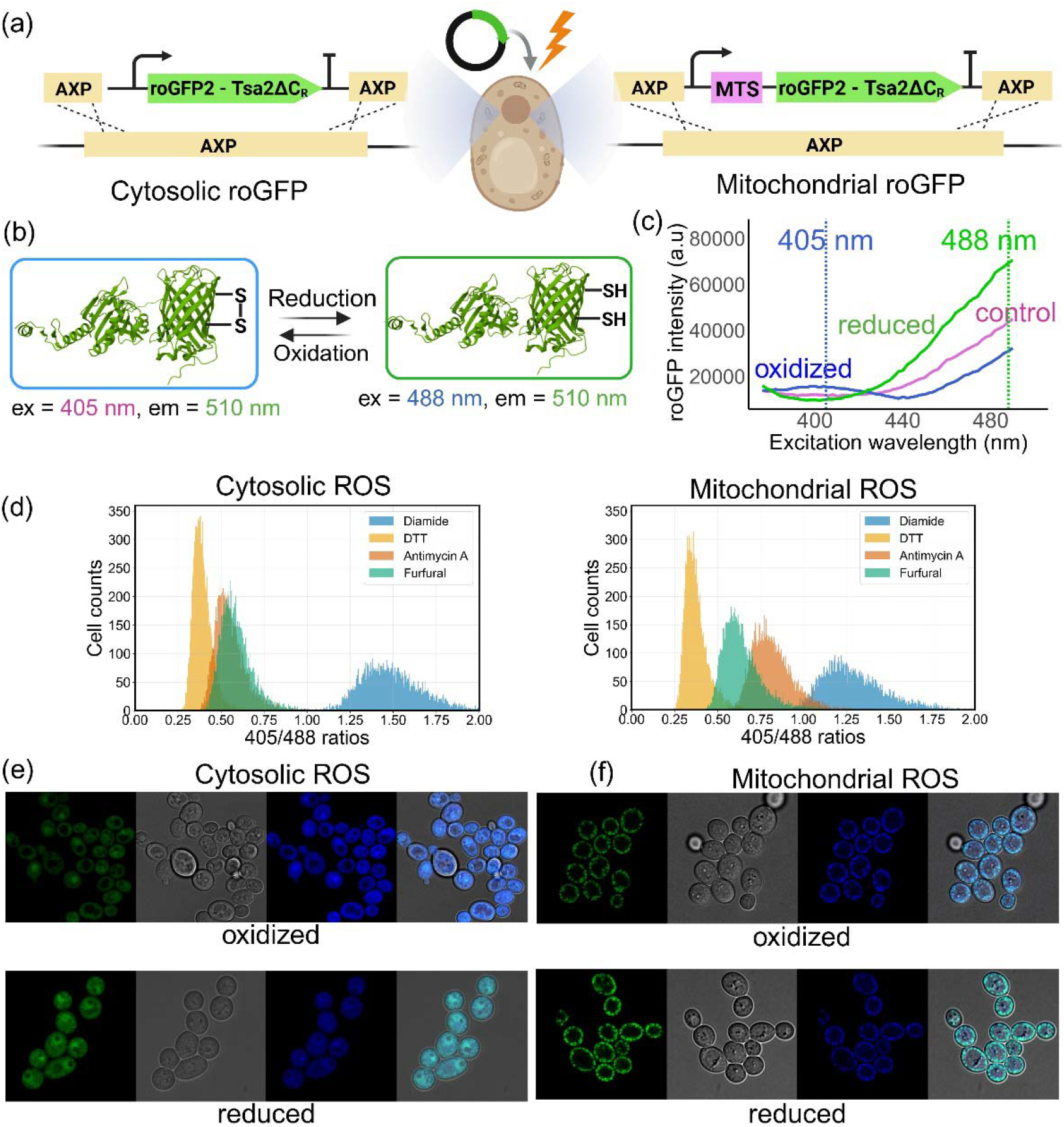
Genetically encoded fluorescent ROS sensor. (a) Construction of cytosolic and mitochondrial roGFP2-Tsa2ΔC_R_ strains via CRISPR-based genomic integration. (b) Schematic of roGFP2-Tsa2ΔC_R_ under oxidized and reduced states. (c) Emission intensity at λ_em_ = 515 nm with excitation wavelengths scanned from 375-488 nm. (d) Flow cytometry histograms of *I*_405_/*I*_488_ ratios for cytosolic and mitochondrial roGFP2-Tsa2ΔC_R_ strains treated with DTT, furfural, AA, and diamide. (e, f) Confocal images of cytosolic (e) and mitochondrial (f) roGFP2-Tsa2ΔC_R_ strains under reduced (DTT) and oxidized (diamide) conditions. Images show 488 nm excitation (green), phase contrast, 405 nm excitation (blue), and merged overlays.

#### 3.1.1. Single-cell level ROS stress measurement

Flow cytometry was used to characterize roGFP2-Tsa2ΔC_R_ responses in individual yeast cells under different redox perturbations. Exponentially growing cells were treated with 20 mM DTT, 20 mM diamide, and varying concentrations of AA, rotenone, furfural, and acetic acid. AA and rotenone inhibit mitochondrial ETC complexes III and I, respectively (Grant et al., 1997; Lai et al., 2008), whereas furfural and acetic acid, used as representative lignocellulosic hydrolysate toxins, are known to impair mitochondrial function and induce oxidative (Kim and Hahn, 2013; Allen et al., 2010; Konzock et al., 2021). As shown in **Fig. 2(d)**, histograms of the *I*_405_/*I*_488_ ratio revealed distinct cytosolic and mitochondrial redox responses. DTT produced the lowest median ratios (~ 0.170 in the cytosol and ~0.164 in mitochondria), while diamide induced strong oxidation with median ratios of 0.804 and 0.675, respectively. Lower mitochondrial ROS levels likely reflect stronger antioxidant buffering by glutathione and peroxiredoxins (Herreto et al., 2008). To enable quantitative comparison across instruments and compartments, OxD values were calculated using DTT- and diamide-treated cells as fully reduced and oxidized references. AA and furfural produced intermediate cytosolic ratios of 0.254 and 0.358, respectively, while mitochondrial AA treatment increased the ratio to 0.425, consistent with direct ETC perturbation. These results demonstrate that roGFP2-Tsa2ΔC_R_ provides a sensitive and quantitative readout of intracellular redox dynamics under diverse chemical stresses.

#### 3.1.2. Imaging intracellular ROS levels

Confocal fluorescence microscopy was used to visualize the subcellular localization and redox response of roGFP2-Tsa2ΔC_R_ in yeast cells. As shown in **Fig. 2 (e)** and **(f)**, cells treated with 20 mM DTT or diamide were imaged under 405 nm and 488 nm excitation together with bright-field imaging. The 405 nm and 488 nm fluorescence channels were pseudocolored blue and green, respectively, and merged with bright-field images. Oxidized cells appeared predominantly blue, whereas reduced cells showed stronger cyan coloration due to enhanced 488 nm excitation. Cytosolic roGFP2-Tsa2ΔC_R_ displayed uniform fluorescence throughout the cell, while mitochondrial roGFP2-Tsa2ΔC_R_ exhibited punctate fluorescence patterns consistent with mitochondrial localization. These results confirm compartment-specific sensor targeting and demonstrate the ability of confocal imaging to resolve local intracellular redox dynamics beyond bulk fluorescence measurements.

### 3.2. Quantification of cellular respiration

Integration of oxygen-sensitive dyes into microwell plates enables real-time monitoring of dissolved oxygen (DO) and cellular OCR (Penso et al., 2021). In this study, PtOEP-doped thin films were incorporated into 96-well plates, as described in section 2.6 (**Fig. 3(a)**). PtOEP fluorescence is inversely proportional to DO concentration because oxygen quenches fluorescence emission. The excitation spectrum of the PtOEP film, measured at 650 nm emission, exhibited peaks at 380 nm and 532 nm (**Fig. 3(b)**). Excitation at 532 nm was selected for respiration assays to minimize UV-induced cellular damage. Air-saturated (~ 0.26 mM DO) and deoxygenated buffers were used as calibration controls. As shown in **Fig. 3(c)**, oxygen depletion increased PtOEP emission intensity by approximately sevenfold at 650 nm under 532 nm excitation. To validate the assay, respiration measurements were performed at cell densities ranging from OD□□□ = 0.05 - 0.4. Cell suspensions (200 μL) were sealed in 96-well plates with soft paraffin to minimize gas exchange, and PtOEP fluorescence was recorded every 2 min for 90 min. As shown in **Fig. 3(d)**, higher cell densities caused faster oxygen depletion and correspondingly greater increases in fluorescence intensity.

**Figure 3.**
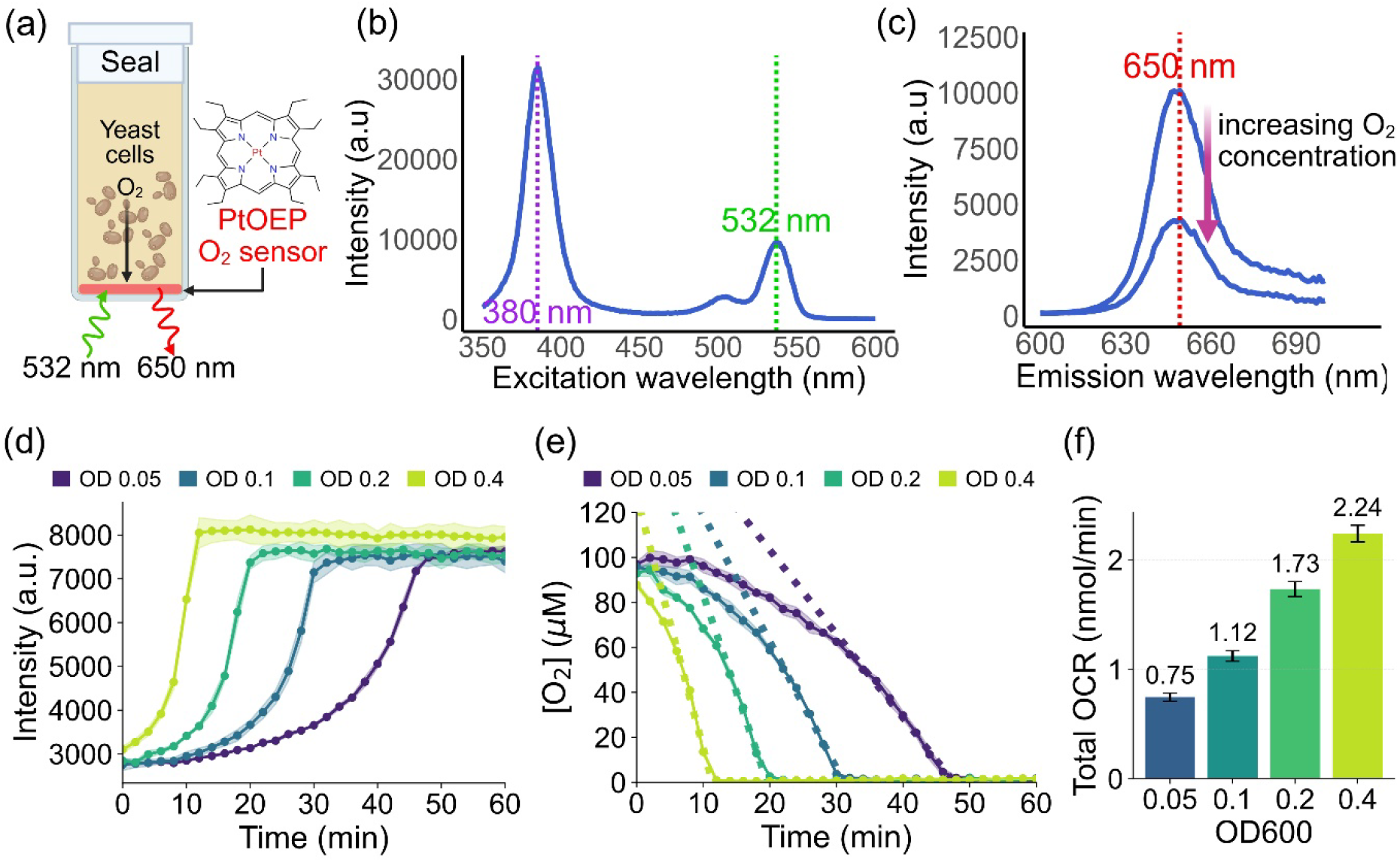
Measurement of yeast respiration. (a) Schematic illustration of the fluorescence-based DO sensor incorporated in 96-well plates. (b) Measured absorption spectrum from the PtOEP incorporated 96-well plate. (c) Measured emission characteristics of the sensor when the PtOEP was excited at 532 nm for different DO levels. (d) Raw time course measurement of PtOEP emission during *Y. lipolytica* respiration. (e) Fitting processed data for the calculation of OCR for *Y. lipolytica* respirations. (f) Calculated OCRs for *Y. lipolytica* at different OD_600_.

To calculate OCRs at different cell densities, raw fluorescence intensity data were converted to dissolved oxygen concentrations using the Stern-Volmer equation, which relates fluorescence quenching to oxygen levels through inversion of the normalized intensity signal, as detailed in section 2.7. The inverted curves in **Fig. 3(e)** depict oxygen concentration as a function of time during cellular respiration. This data were subsequently fitted with a linear regression model in a dynamic window, with the slope corresponding to the rate of cellular oxygen consumption. The OCRs obtained for cultures with OD_600_ values of 0.05, 0.1, 0.2, and 0.4 were 0.75, 1.12, 1.73, and 2.24 nmol/min, respectively, demonstrating a strong positive correlation between biomass concentration and respiratory activity. Additionally, the oxygen consumption profile of individual wells remained independent, with no observable crosstalk between the wells. These results validate the PtOEP-based microwell assay as a robust and high - throughput method for quantifying cell respiration dynamics across varying cell densities.

### 3.3. Quantitative assessment of yeast stains with different metabolic profiles

Different yeast species exhibit distinct respiratory strategies that produce characteristic metabolic and intracellular redox states. Using the integrated sensing platform, we compared intracellular ROS levels and oxygen consumption rates across yeast strains and cultivation conditions representing different respiratory regimes. Glucose-grown Crabtree-positive *S. cerevisiae* served as fermentative, low-respiration state, whereas glycerol-grown *S. cerevisiae* represented enhanced mitochondrial respiration. In contrast, the obligate aerobic and Crabtree-negative yeast *Y. lipolytica* exhibited intrinsically high respiratory activity during glucose cultivation. Comparing these physiological states enabled quantitative analysis of the relationship between respiration and intracellular redox balance under fermentative and respiratory metabolism.

#### 3.3.1 Basal level of yeast cell respiration rates and ROS levels

Basal respiratory activity and intracellular oxidative stress were quantified under unperturbed growth conditions to establish reference physiological states for each yeast system. Glucose- and glycerol-grown *S. cerevisiae* were used to represent fermentative and respiratory metabolism, respectively, while glucose-grown *Y. lipolytica* represented a highly respiratory state (Imura et al., 2020). All measurements were performed at a normalized cell density (OD□□□ = 0.2) to minimize biomass-related variability. As shown in **Fig. 4(d)**, glucose-grown *S. cerevisiae* exhibited the lowest OCR at 4.84 nmol min□^1^ OD□□□□^1^, consistent with fermentative metabolism. Growth on glycerol increased OCR to 11.85 nmol min□^1^ OD□□□□^1^ (~2.5-fold increase), reflecting enhanced mitochondrial respiration. In contrast, glucose-grown *Y. lipolytica* displayed the highest OCR at 33 nmol min□^1^ OD□□□□^1^, approximately 6.8-fold and 3-fold higher than glucose- and glycerol-grown *S. cerevisiae*, respectively.

**Figure 4.**
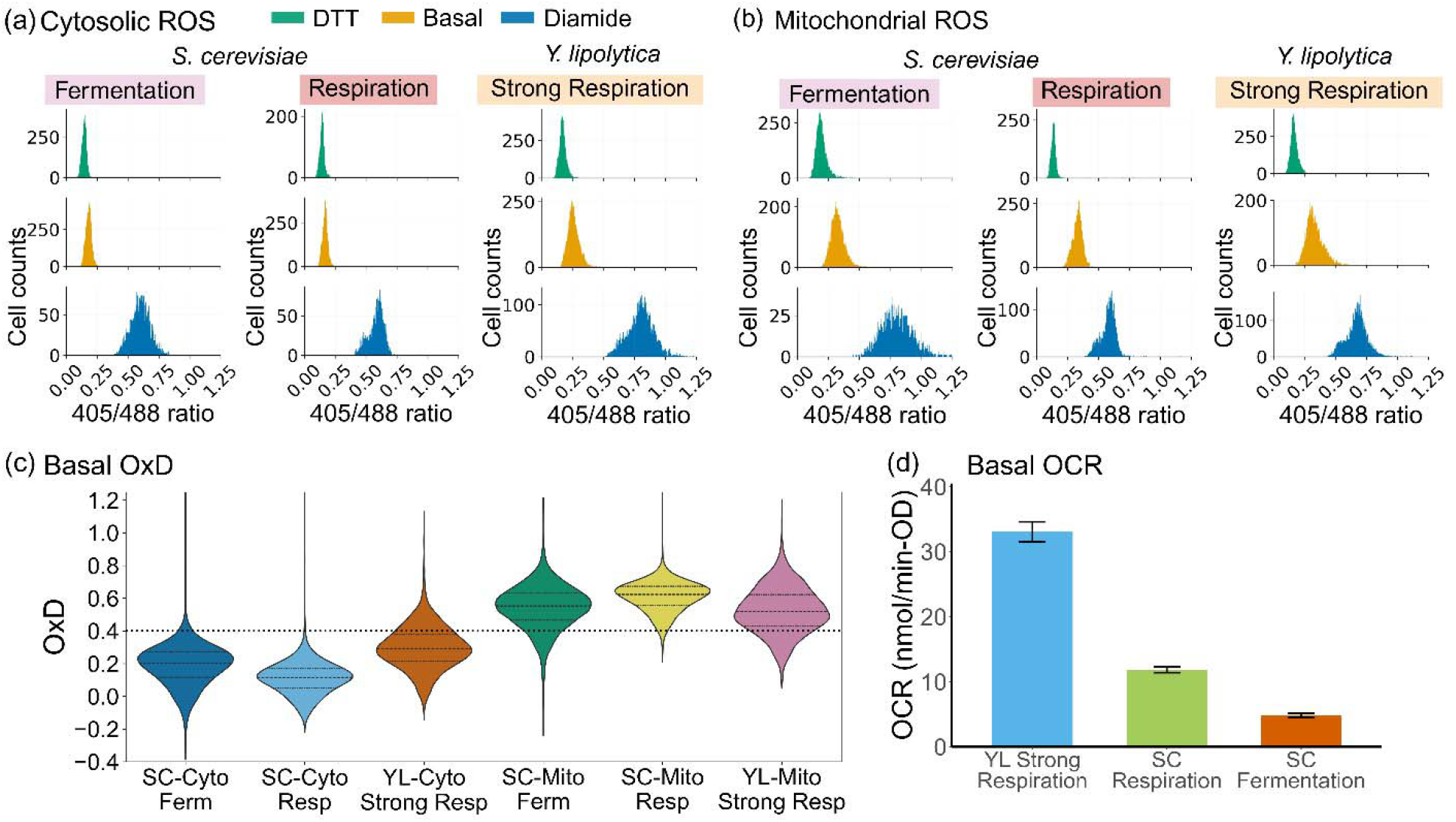
Basal respiration and ROS levels of *S. cerevisiae* and *Y. lipolytica*. (a, b) *I*_405_/*I*_488_ ratios in the cytosol (a) and mitochondria (b) of glucose-grown and glycerol-grown *S. cerevisiae*, and glucose-grown *Y. lipolytica*, representing fermentative, respiratory, and strong respiratory states, respectively. Measurements were performed at OD□□□ = 0.2. (c) Basal OxD levels of the three cultures. (d) Calculated basal OCR under different metabolic conditions.

Intracellular ROS levels measured using the roGFP2-Tsa2ΔC_R_ sensor are shown in **Fig. 4(c)**. Glucose-grown *S. cerevisiae* exhibited the lowest basal oxidative stress, with median OxD values of 0.204 in the cytosol and 0.553 in mitochondria. Glycerol-grown *S. cerevisiae* showed lower cytosolic OxD of 0.110 but increased mitochondrial OxD of 0.623, consistent with enhanced respiratory metabolism. *Y. lipolytica* displayed OxD values of 0.293 in the cytosol and 0.521 in mitochondria, reflecting sustained respiratory activity. Across all growth conditions, the cytosol remained more reduced than mitochondria, consistent with known compartment-specific redox properties (López-Mirabal and Winther, 2008; Colombini, 2004). The distinct OxD profiles further indicate that cytosolic and mitochondrial redox homeostasis respond differently depending on metabolic state, carbon source, and species-specific respiratory pathways (De Cubas et al., 2021; Mishina et al., 2019). Together, these results demonstrate a quantitative relationship between basal respiration and intracellular redox state across yeast strains with distinct metabolic strategies.

#### 3.3.2 Chemical-induced ROS imbalance and OCR inhibition of S. cerevisiae

To investigate how metabolic state influences redox sensitivity to respiratory perturbations, exponentially growing *S. cerevisiae* cells cultured in glucose or glycerol were exposed to chemical stressors. Antimycin A (AA), an ETC complex III inhibitor, was applied at increasing concentrations to perturb mitochondrial respiration, as well as increasing concentrations of furfural and acetic acid, known to impair mitochondrial function and disrupt metabolism and redox balance. Changes in OCR and intracellular redox state were quantified relative to untreated controls, enabling systematic comparison of stress responses under fermentative and respiratory metabolic conditions.

As shown in **Fig. 5(a)**, antimycin A induced a concentration-dependent decline in OCR in *S. cerevisiae* under both metabolic conditions. In glucose-grown fermenting cells, OCR decreased by 40% from 4.83 to 2.86 nmol min□^1^ OD□□□□^1^ at 1.25 µM AA, whereas glycerol-grown respiring cells showed greater sensitivity, with an 85% reduction from 11.85 to 1.74 nmol min□^1^ OD□□□□^1^. Furfural also caused concentration-dependent OCR suppression, with stronger effects in respiring cells. At 10 mM furfural, OCR decreased by 13% and 35% in glucose- and glycerol-grown cultures, respectively, indicating greater vulnerability of respiration-dependent cells to mitochondrial stress (Barros et al., 2004; Li et al., 2022). Acetic acid produced distinct responses depending on metabolic state. In glucose-grown cells, OCR increased across 2.5 - 40 mM, rising by 35 - 42%, consistent with its utilization as an additional carbon source (Chaves et al., 2021). In contrast, glycerol-grown cells showed increased OCR at low concentrations (2.5 - 5 mM) but progressive suppression at higher concentrations, with OCR decreasing by 55% at 40 mM. These results demonstrate that metabolic state strongly influences physiological responses to respiratory and chemical stressors. At 80 mM, acetic acid was acutely toxic under all conditions.

**Figure 5.**
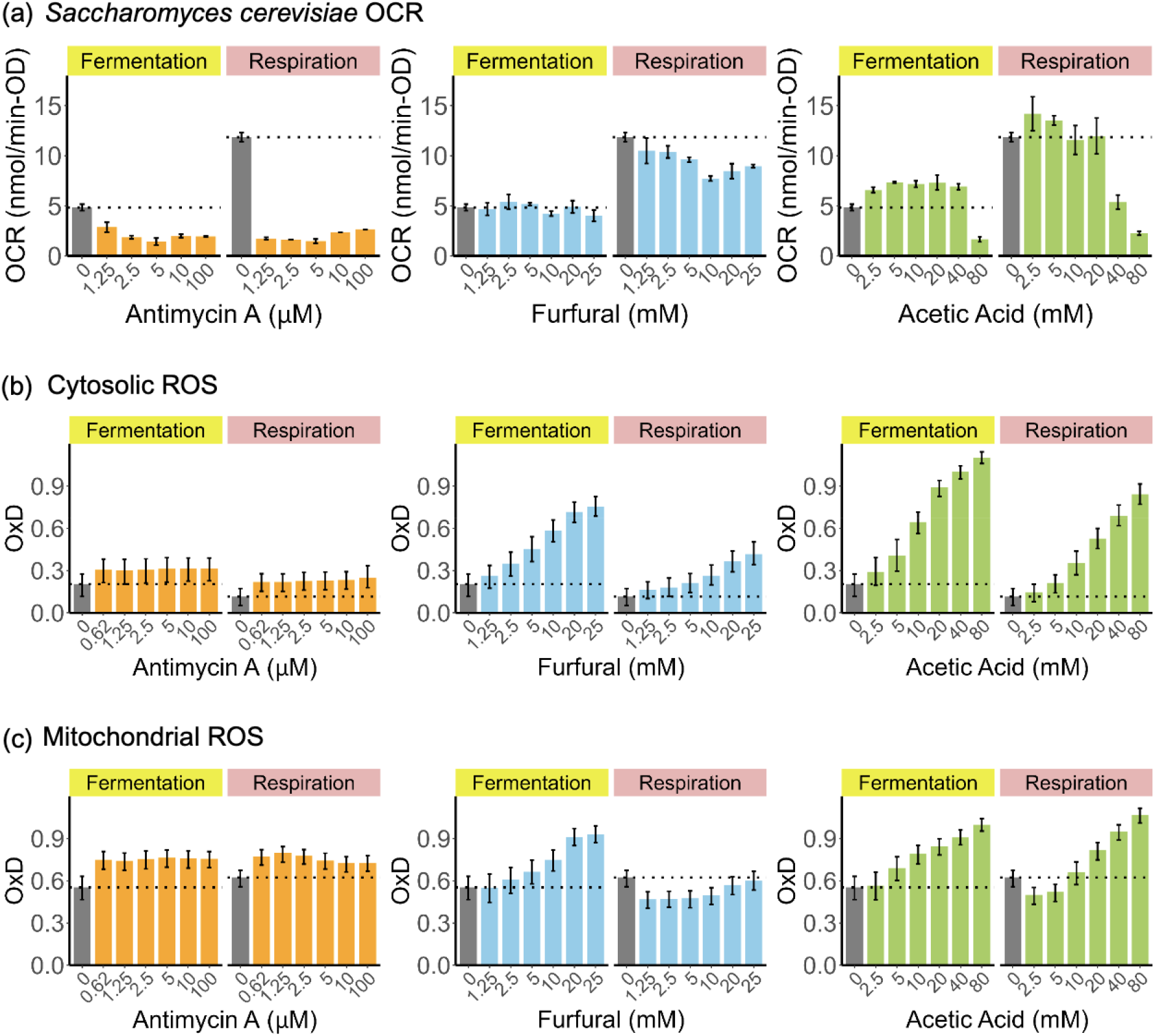
Chemical-induced ROS imbalance in *S. cerevisiae*. (a) OCR of *S. cerevisiae* treated with different concentrations of AA, furfural, and acetic acid, compared with untreated controls in isosmotic buffer. (b, c) Cytosolic (b) and mitochondrial (c) OxD responses under the same treatment conditions.

Intracellular ROS responses measured using the cytosolic roGFP2-Tsa2ΔC_R_ sensor are shown in **Fig. 5(b)**. In fermenting *S. cerevisiae*, cytosolic OxD increased across a broad range of antimycin A concentrations, rising from a basal value of 0.203 to 0.301 (50% increase) at 1.25 µM AA. Respiring cells exhibited a lower basal OxD of 0.115 but showed a stronger relative response to ETC inhibition, increasing to 0.218 (90% increase) at 1.25 µM AA. Furfural and acetic acid induced larger concentration-dependent increases in cytosolic OxD than AA under both metabolic conditions. In fermenting cells, furfural increased OxD to 0.584 (~ 2.8-fold increase) at 10 mM, while acetic acid elevated OxD to 1.000 (~ 4.9-fold increase) at 40 mM. Respiring cells maintained lower overall OxD levels but still showed pronounced oxidative responses, with acetic acid increasing OxD from 0.115 to 0.691 (~ 6-fold increase) at 40 mM. These results indicate that although respiring *S. cerevisiae* maintains lower basal cytosolic ROS, mitochondrial perturbation strongly enhances oxidative stress.

Basal mitochondrial OxD levels were higher than cytosolic values but showed smaller responses to external perturbations (**Fig. 5(c)**). In fermenting *S. cerevisiae*, mitochondrial OxD increased from 0.553 to 0.741 (34% increase) following 1.25 µM AA treatment and remained elevated across all AA concentrations tested. In respiring cells, mitochondrial OxD increased from 0.623 to 0.796 (27% increase) at 1.25 µM AA, then gradually declined at higher concentrations while remaining above basal levels. Furfural induced concentration-dependent mitochondrial oxidation in fermenting cells, increasing OxD by 35% to 0.749 at 10 mM. In contrast, respiring cells maintained mitochondrial OxD below basal levels across all furfural concentrations, suggesting stronger redox buffering capacity. Acetic acid produced concentration-dependent mitochondrial oxidation under both metabolic conditions. In fermenting cells, OxD increased to 0.908 (64% increase) at 40 mM, while in respiring cells low concentrations reduced OxD, but high concentrations increased OxD to 0.948 (52% increase).

Comparison of mitochondrial and cytosolic OxD values showed that mitochondria maintain higher basal redox levels but exhibit smaller relative changes under external perturbations. Fermenting cells displayed higher overall OxD, whereas respiring cells showed smaller and more tightly regulated redox shifts. The reduced mitochondrial OxD fluctuations in glycerol-grown cells likely reflect enhanced respiratory flux and antioxidant capacity that more effectively buffer transient oxidative stress.

#### 3.3.3 Chemical stresses induced ROS imbalance and respiration response in Y. lipolytica

To assess strain-specific differences in redox and respiratory responses to mitochondrial perturbation, exponentially growing *Y*.□*lipolytica* cells were cultured and exposed to the ETC inhibitors and toxic compounds, furfural and acetic acid, at increasing concentrations to evaluate their impact on this aerobic yeast. As shown in **Fig. 6(a)**, *Y. lipolytica* exhibited minimal OCR suppression in response to ETC inhibitors. Basal OCR in untreated cells was 33 nmol min□^1^ OD□□□□^1^ and remained nearly unchanged across 1.25–10 µM rotenone, with only a 25% reduction at 100 µM. Unexpectedly, antimycin A (AA) stimulated respiration, increasing OCR by 19% to 39.42 nmol min□^1^ OD□□□□^1^ at 1.25 µM before gradually declining at higher concentrations while remaining near or above basal levels. These responses contrast with those observed in *S. cerevisiae*, indicating greater respiratory resilience in *Y. lipolytica*. Similar to respiring *S. cerevisiae*, furfural caused concentration-dependent OCR suppression, reducing OCR by 66% at 10 mM. In contrast, acetic acid produced a biphasic response, increasing OCR across 2.5–40 mM and peaking at 45.25 nmol min□^1^ OD□□□□^1^ (36% increase) at 40 mM, followed by a sharp 80% decrease at 80 mM. The mechanism underlying this biphasic OCR response remains unclear.

**Figure 6.**
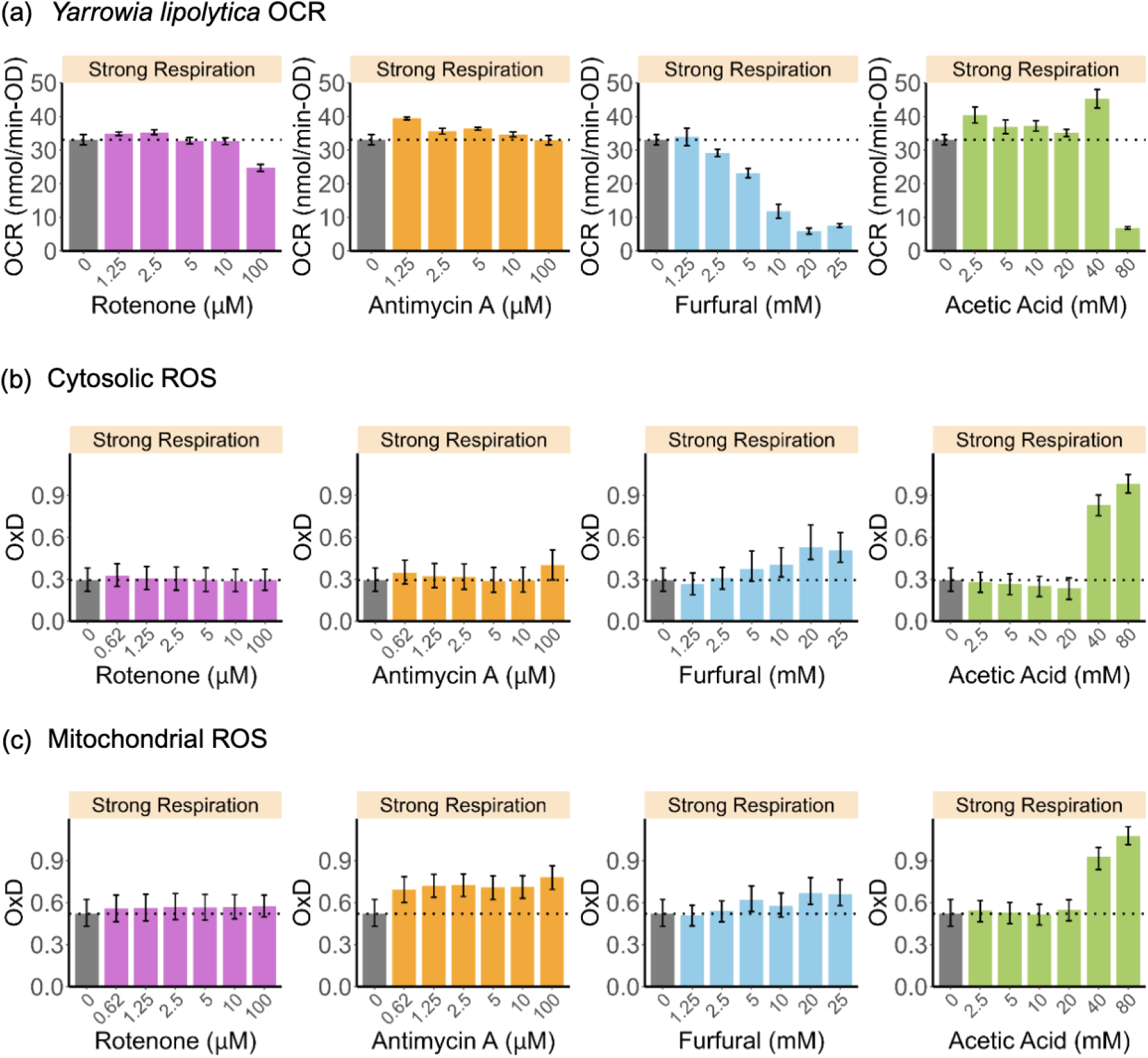
Chemical-induced ROS imbalance in *Y. lipolytica*. (a) OCR of *Y. lipolytica* treated with different concentrations of rotenone, AA, furfural, and acetic acid, compared with untreated controls in isosmotic buffer. (b, c) Cytosolic (b) and mitochondrial (c) OxD responses under the same treatment conditions.

Intracellular ROS responses measured using the cytosolic roGFP2-Tsa2ΔC_R_ sensor are shown in **Fig. 6(b)**. In *Y. lipolytica*, cytosolic OxD remained largely stable across rotenone (0.625 - 100 µM) and antimycin A (0.625 - 10 µM) treatments, with only 100 µM AA increasing OxD by 36% to 0.401. Furfural induced a concentration-dependent increase in OxD, rising from near basal levels at 1.25 mM to 0.532 (81% increase) at 20 mM. Acetic acid produced a biphasic response similar to OCR behavior, with OxD decreasing slightly at 2.5 - 20 mM before sharply increasing to 0.979 (~3.3-fold above basal) at 80 mM. Mitochondrial redox responses (**Fig. 6(c)**) showed similar trends. Rotenone caused minimal changes, with OxD increasing only 10% at 100 µM. In contrast, AA produced a concentration-dependent increase in mitochondrial OxD, reaching 0.781 (50% increase) at 100 µM. Furfural also progressively elevated mitochondrial OxD to 0.669 (28% increase) at 20 mM. Acetic acid maintained near-basal OxD at 2.5 - 20 mM but increased mitochondrial OxD sharply at higher concentrations, reaching 0.929 at 40 mM and 1.076 (~2-fold increase) at 80 mM.

Overall, these results demonstrate that chemical inhibition of mitochondrial respiration induces dose-dependent OCR suppression and compartment-specific ROS accumulation in both *S. cerevisiae* and *Y. lipolytica*. In *S. cerevisiae*, fermenting cells exhibited lower respiratory inhibition but greater oxidative stress than respiring cells, highlighting the dependence of redox vulnerability on metabolic state. In contrast, *Y. lipolytica* maintained more stable respiration and redox homeostasis under both mitochondrial and chemical stress, exhibiting attenuated OCR suppression and reduced ROS accumulation. This enhanced resilience is consistent with its obligate respiratory metabolism and the presence of auxiliary ETC components, including external NADH dehydrogenase and alternative oxidase (AOX), which provide bypass electron transfer pathways independent of the canonical Complex I-IV sequence (Medentsev et al., 2002, Li et al., 2003; Guerrero-Castillo et al., 2009 and 2012]. Together, the integrated OCR and roGFP measurements establish a quantitative link between ETC perturbation, metabolic state, and intracellular redox dynamics, revealing strain-specific strategies for maintaining oxidative balance.

## 4. Conclusion

In this study, we developed and validated an integrated dual-sensing platform combining the genetically encoded roGFP2-Tsa2ΔC_R_ redox biosensor with a PtOEP-based OCR assay to quantitatively link intracellular redox dynamics with extracellular respiratory activity in yeast. This approach enabled simultaneous, compartment-resolved measurement of ROS-dependent OxD and whole-cell oxygen consumption across distinct species and metabolic states, providing a quantitative view of cellular fitness under physiological and chemically perturbed conditions. Our results further demonstrate that mitochondria and the cytosol maintain distinct redox states with different sensitivities to external perturbations.

Under basal growth conditions, fermentation-dominant *S. cerevisiae* grown in glucose exhibited low OCR, whereas glycerol-grown *S. cerevisiae* and aerobic *Y. lipolytica* showed progressively higher respiratory activity. Increased OCR correlated with elevated mitochondrial ROS but lower cytosolic ROS, indicating compartmentalized oxidative burden during respiratory metabolism. Under chemical perturbation, metabolic state strongly influenced redox sensitivity in *S. cerevisiae*, where respiring cells showed greater OCR suppression and ROS accumulation following ETC inhibition and lignocellulosic toxin exposure. Furfural consistently impaired respiration and increased OxD, while acetic acid produced biphasic responses, enhancing respiration at low concentrations but causing severe oxidative stress at higher doses. In contrast, *Y. lipolytica* maintained greater respiratory robustness and redox stability, exhibiting only modest OCR and OxD changes even under high rotenone and antimycin A concentrations.

Together, the combined OCR and roGFP2-Tsa2ΔC_R_ measurements reveal species- and pathway-specific strategies for maintaining oxidative balance. By coupling intracellular redox sensing with optical respiration measurements, this platform provides a scalable tool for linking redox state, energy metabolism, and cellular fitness. Such integrated monitoring could support strain optimization, stress characterization, and real-time bioprocess monitoring in microbial biomanufacturing. Future integration with complementary sensors for pH, metabolites, or ATP may further expand its utility for multidimensional characterization of host physiology and process performance.

## Acknowledgements

This research was partially supported by the National Institutes of Health under Award No. R35GM143048 and the National Science Foundation under Award No. 2533046. The content is solely the responsibility of the authors and does not necessarily represent the official views of the National Institutes of Health or the National Science Foundation. Some parts of the figures were made in BioRender (https://BioRender.com)

